# Macrophage-induced rifampin tolerance across Mycobacterium tuberculosis lineages is Rv1258c-dependent

**DOI:** 10.1101/383109

**Authors:** Kristin N. Adams, Amit Kumar Verma, Radha Gopalaswamy, Harresh Adikesavalu, Dinesh Kumar Singhal, Srikanth Tripathy, Uma Devi Ranganathan, David R. Sherman, Kevin B. Urdahl, Lalita Ramakrishnan, Rafael E. Hernandez

## Abstract

The Mycobacterium tuberculosis (Mtb) Lineage 4 strains CDC1551 and H37Rv develop tolerance to multiple antibiotics upon macrophage residence. Genetic mutation of the efflux pump Rv1258c in CDC1551 abolishes rifampin tolerance but not isoniazid tolerance. Here we show that clinical isolates from the other predominant Mtb lineages developed macrophage-induced isoniazid tolerance. Furthermore, all lineages developed rifampin tolerance except Lineage 2 Beijing strains, which are natural Rv1258c mutants. Thus macrophage-induced antibiotic tolerance is featured across the majority of Mtb lineages. Our findings further link Rv1258c to rifampin tolerance among clinical isolates.

## BACKGROUND

Mycobacterium tuberculosis (Mtb) enters host macrophages shortly after infection and resides within granulomas, organized macrophage aggregates, for much of its life cycle [1]. Previously we and others showed that Mtb develops tolerance to multiple first and second line anti-tubercular drugs soon after infecting macrophages [2–4]. Moreover, we found that macrophage-induced tolerance to rifampin is mediated via Tap (Rv1258c), a major facilitator superfamily (MFS) efflux pump [2]. Rv1258c expression is induced when bacteria reside within cultured human macrophages [5] as well as in bacteria in sputum of TB patients undergoing treatment with a rifampin-containing regimen [6]. These observations suggest that macrophage induced tolerance to rifampin mediated by Rv1258c may contribute to drug tolerance observed in patients.

Based on genomic differences, Mtb is broadly categorized into multiple lineages associated with distinct phenotypes with regard to mutability, drug sensitivity, immunogenicity and virulence [7]. The vast majority of TB worldwide (over 90%) is caused by Mtb lineages 1 (Indo-oceanic), 2 (East Asian), 3(East African-Indian), and 4 (Euro-American)[7](Supplementary Figure 1). Our prior observations of macrophage induced tolerance were made in H37Rv and CDC1551, both of which represent lineage 4 strains. While lineage 4 is perhaps the most widely distributed geographically, it accounts for only approximately 11% of the global TB burden [7] (Supplementary Figure 1B). Therefore, we sought to determine whether macrophage-induced tolerance to isoniazid and rifampin is common across the other three lineages that are predominant in high TB-burden areas [7]. In addition, because the Beijing subgroup of lineage 2 strains harbor an inactivating frameshift mutation in Rv1258c [8], we were interested to see whether they develop rifampin tolerance or not. Furthermore, Rv1258c also facilitates bacterial growth within macrophages[2,9], so we assessed both Beijing and non-Beijing strains for growth within macrophages.

## METHODS

### Bacterial strains

The sources and antibiotic susceptibilities of the strains used are detailed in Supplementary Table 1. Bacteria were grown to mid log-phase in Middlebrook 7H9 medium (Becton Dickinson) with 0.05% Tween-80 and albumin, dextrose, catalase (Middlebrook ADC Enrichment, Becton Dickson) prior to infection.

**Figure 1.**
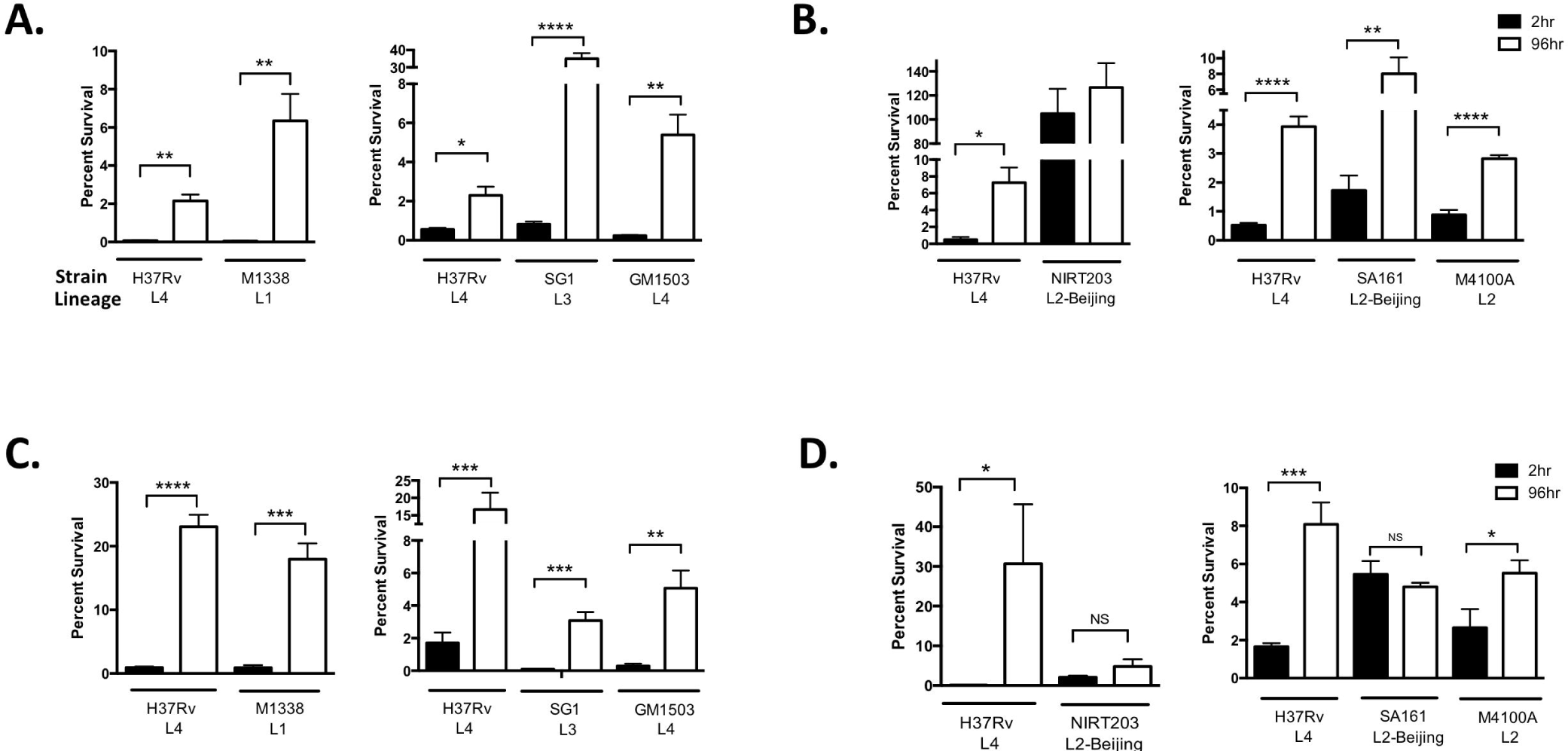
Macrophage induced tolerance to rifampin is common across clinical lineages of M. tuberculosis. A-D, THP-1 macrophages were infected with H37Rv (reference strain) or clinical strains as indicated and lysed at 2 hours (black bars) or 96 hours (white bars) post-infection. The released bacteria were treated for an additional 48 hours with 0.6 μg/ml isoniazid (A and B) or 1 μg /ml rifampicin (C and D) prior to enumeration of colony-forming units (CFU). Results (A, C and D) are representative of three independent experiments and (B) is representative of two experiments. Error bars represent standard deviation. Significance testing was performed using T-test.

#### Macrophage Growth and Infection

THP-1 cells (ATCC) were grown in RPMI 1640, supplemented with 10% FBS and 2mM L-glutamine (Sigma) in 37°C incubator with 5% CO2. 5×10^5^THP-1 cells were differentiated into 24-well plates with 100 nM phorbol 12-myristate 13-acetate (Sigma) for 48 hours and allowed to recover for 24 hours prior to infection. The differentiated cells were infected at a multiplicity of infection (MOI) of 1 for 2 hours. Cells were washed with media and 6 μg/ml streptomycin (Sigma) was added to the media for the remainder of the intracellular growth; this was defined as the start of infection. Media were changed every 48 hours. For intracellular growth inhibition assays, verapamil HCl (40 μg/ml) (Sigma) was added to the media 48 hours post-infection and streptomycin was omitted.

#### Macrophage-induced tolerance assay

The work flow for this assay is depicted in Supplementary Figure 2. Briefly, THP1 cells were infected as above and were lysed 2 or 96 hours post-infection to release the bacteria as follows: Cells were washed briefly once with PBS and then with water. Cells were then incubated with 100 μl of water per well at 37°C for 15 minutes. Then 900 μl of 7H9 medium (supplemented with Middlebrook ADC and 0.05% Tween-80) was added and the wells bottoms scraped with a pipette tip to ensure complete macrophage lysis, which was microscopically confirmed. Serial dilutions of 150 μL of cell lysates were made in PBS and plated on 7H10 agar (Becton Dickson) to obtain the initial colony forming units (CFU). To measure antibiotic killing, 500 μL of cell lysate was treated with the indicated antibiotic (rifampin 1 μg/ml or isoniazid 0.6 μg/mL, Sigma) for 48 hours at 37C° before making serial dilutions and plating on 7H10 agar. Percent survival was determined by dividing the post-treatment CFU by the pre-treatment CFU.

**Figure 2.**
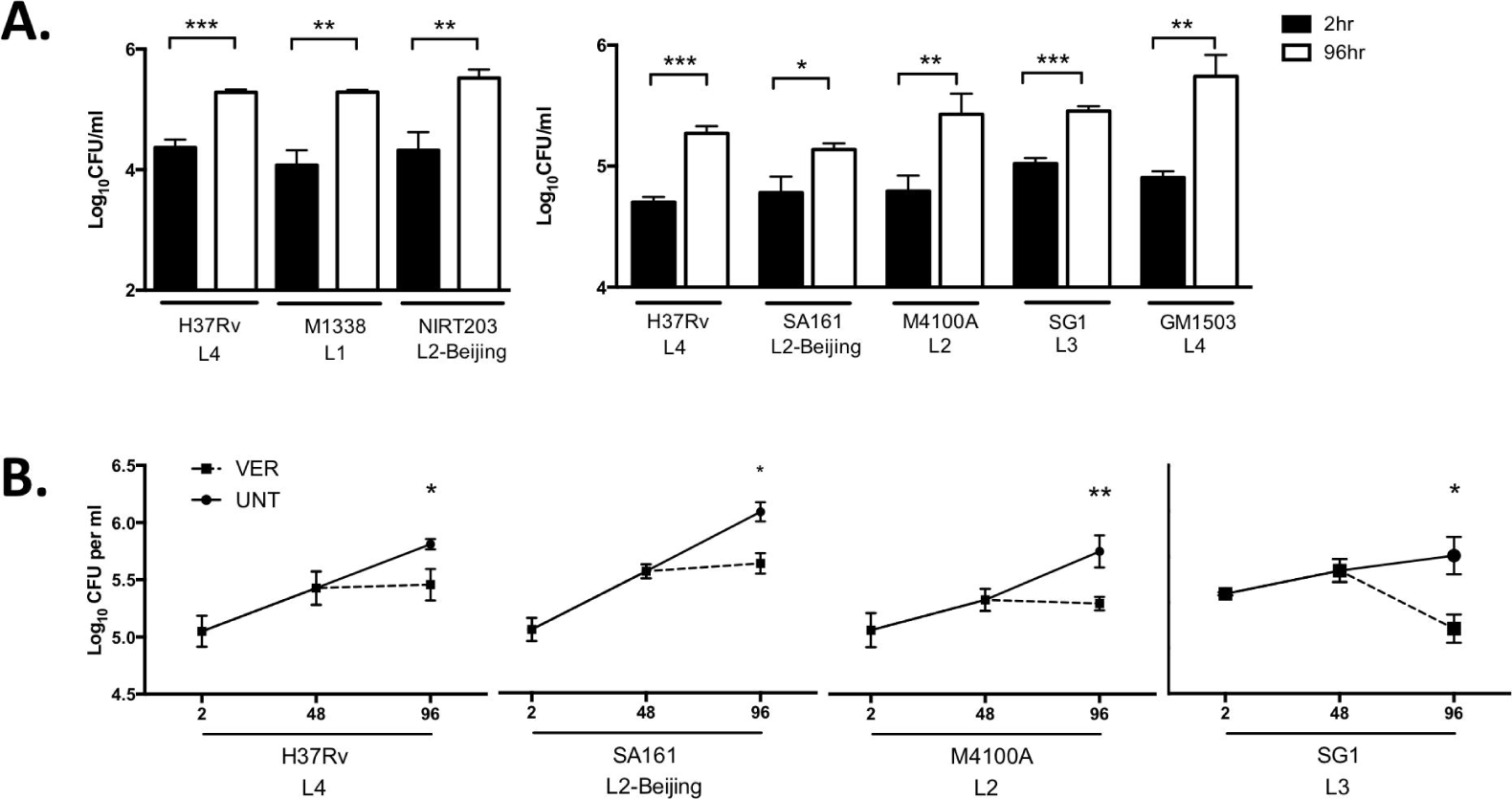
Beijing lineage strains of M. tuberculosis are not compromised for early macrophage growth and are susceptible to intracellular verapamil treatment. A. THP-1 macrophages were infected with H37Rv or clinical strains of M. tuberculosis (MTB) as indicated and lysed at 2 hours (black bars) or 96 hours (white bars) and CFU enumerated at each time-point. B. THP-1 macrophages were infected with MTB strains H37Rv, SA161, M4100A and SG1 for 48 hours and subsequently left untreated or treated for an additional 48 hours with 40 μg/ml verapamil (VER) prior to lysis and enumeration of CFU. Results (A) are representative of at least three independent experiments and (B) is representative of at least two experiments. Error bars represent standard deviation. Significance testing performed using T-test.

#### Intracellular growth assay

Infected cells were washed twice with PBS and incubated with 100 μl 0.1% Triton X-100 for 10 minutes. Then 900 μl of PBS was added and the wells scraped with a pipette tip. Dilutions of cell lysates were plated on 7H10 agar as above.

#### Statistical analyses

GraphPad Prism 6.0 was used for statistical analyses. Means were compared via statistical tests indicated in the figure legends. P values are abbreviated in figures as follows: *, P<0.05; **, P<0.01; *** P<0.001, **** P<0.0001. The number of times experiments were repeated is indicated in each of the figure legends, with results from one representative experiment shown in each figure.

## RESULTS

### Macrophage-induced antibiotic tolerance occurs across predominant Mycobacterium tuberculosis lineages

Working at two sites, Seattle Children’s Research Institute (Seattle, USA) and the National Institute for Research in Tuberculosis (Chennai, India), we used a panel of Mtb strains representing lineages 1-4 assembled from previously published strains (Seattle site) and from recent clinical isolates (Chennai site) (Supplementary Table 1). All strains were confirmed to be susceptible to both isoniazid and rifampin, except strain NIRT203 which was resistant to isoniazid (Supplementary Table 1). The lineage 2 Beijing strains were confirmed to harbor the frameshift mutation in Rv1258c, and this mutation was absent in all other strains, including the lineage 2 non-Beijing isolate M4100A (Supplementary Table 1).

To assess development of macrophage-induced antibiotic tolerance, THP-1 macrophages were infected with the clinical MTB strains and then lysed at 2 and 96 hours post-infection. Macrophage lysates were treated with antibiotics for 48 hours prior to enumerating the bacteria (Supplemental Figure 2A). Macrophage-induced tolerance was defined as a significant (p ≤ 0.05) increase in the fraction of bacteria surviving antibiotic treatment between the 2 hour and 96 hour time points. All of the isoniazid susceptible strains developed macrophage-induced tolerance to isoniazid (Figure 1A and 1B), while NIRT203 was resistant to killing by isoniazid as expected. Strains from lineages 1, 3 and 4 developed tolerance to rifampin (Figure 1C). The capacity of lineage 2 Mtb strains to develop tolerance to rifampin correlated with whether the Rv1258c gene remained intact. Both lineage 2 Beijing strains failed to develop rifampin tolerance (Figure 1D). In contrast, M4100A, a non-Beijing lineage 2 strain that does not harbor the inactivating mutation in Rv1258c, was rifampin tolerant (Figure 1D).

## Lineage 2 Beijing strains grow normally in macrophages

For the CDC1551 strain, Rv1258c mutants not only fail to develop macrophage-induced rifampin tolerance, but also are defective for early growth in macrophages [2,9]. However, Beijing strains do not exhibit a macrophage growth defect; indeed, many of them grow more rapidly in macrophages than non-Beijing isolates [10]. This may be one of the reasons that some Beijing strains have been found to be hypervirulent in animal infection models and are spreading globally [7,11].

When we tested the two Beijing strains in our panel for their ability to grow in macrophages, we found that neither manifested an intramacrophage growth defect, confirming prior findings (Figure 2A). Thus, these Beijing strains have evolved compensatory mechanisms that allow them to grow in macrophages. Additional mechanisms that include host immune dysregulation have been invoked to further render them hypervirulent [7]. Indeed, when we assessed the SA161 Beijing strain in a mouse aerosol infection model, we found it to be hypervirulent. Transient early increased bacterial burdens compared to H37Rv were associated with early lethality (Supplementary Figure 3). Together these findings confirmed that SA161 has not only compensated for any macrophage growth defect due to the loss of Rv1258c but has further evolved additional mechanisms that renders it hypervirulent. Because multiple efflux pumps are reported to be upregulated in Beijing strains [12], we considered the possibility that the mechanisms that compensated for its early growth in macrophages might include the induction of other efflux pumps. Consistent with this possibility, we found that the bacterial efflux pump inhibitor verapamil inhibited SA161’s intramacrophage growth similarly to strains from other lineages [13](Figure 2B).

## DISCUSSION

Our earlier studies showed that macrophage-induced drug tolerance is a potential contributor to the slow response of Mtb to antimicrobial treatment [2,3]. However, the findings were limited to two laboratory strains belonging to a single Mtb lineage. This work shows that macrophage-induced drug tolerance is a feature of the other three predominant Mtb lineages as well. Moreover, the finding that the Beijing strains lack rifampin tolerance while retaining isoniazid tolerance corroborates our previous findings linking the efflux pump to the development of macrophage-induced rifampin tolerance [2].

Our finding that the Beijing strains fail to develop macrophage-induced rifampin tolerance might be seen as presenting a potential quandary given that Beijing lineage TB is more likely to relapse after standard rifampin-containing regimens [14]. However, this increased propensity to relapse may simply be due to the compensated growth in macrophages and hypervirulence traits of the Beijing lineage, as we have shown here for SA161. Our finding that verapamil inhibits intramacrophage growth of Beijing strains suggests that adjunctive therapy with it or other efflux pump inhibitors may be useful for the treatment of Beijing strain-caused tuberculosis to at least partially overcome these compensatory virulence mechanisms.

## FOOTNOTES

### Author Contributions

K.N.A., A.K.V., K.U.D., L.R. and R.E.H. designed experiments. K.N.A., A.K.V., R.G., H.A., D.K.S. and R.E.H. performed experiments. K.N.A., A.K.V., D.R.S., K.B.U., U.D.R., L.R. and R.E.H. analyzed and interpreted data. U.D.R., D.R.S. and S.T. provided project administration and supervision. K.N.A., L.R. and R.E.H. prepared figures and wrote the manuscript. All authors reviewed the manuscript. L.R. conceived the project.

## Acknowledgements

We thank Ian Orme for providing the SA161 strain, Shiva Kumar for providing the M1338 and NITR203 strains, Louie Galitan for technical assistance and Paul Edelstein for providing input on the manuscript.

## Prior presentation

Portions of this work were previously presented at the Tuberculosis Drug Discovery and Development Gordon Research Conference at Lucca, Italy (June 2017) and at the PacTB Symposium in Seattle, WA (March 2018).

